# Inferring expressed genes by whole-genome sequencing of plasma DNA

**DOI:** 10.1101/049478

**Authors:** Peter Ulz, Gerhard G. Thallinger, Martina Auer, Ricarda Graf, Karl Kashofer, Stephan W. Jahn, Luca Abete, Gunda Pristauz, Edgar Petru, Jochen B. Geigl, Ellen Heitzer, Michael R. Speicher

## Abstract

The analysis of cell-free DNA (cfDNA) in plasma represents a rapidly advancing field in medicine. cfDNA consists predominantly of nucleosome-protected DNA shed into the bloodstream by cells undergoing apoptosis. We performed whole-genome sequencing (WGS) of plasma DNA and identified two discrete regions at transcription start sites (TSS) where the nucleosome occupancy results in different read-depth coverage patterns in expressed and silent genes. By employing machine learning for gene classification, we found that the plasma DNA read depth patterns from healthy donors reflected the expression signature of hematopoietic cells. In cancer patients with metastatic disease, we were able to classify expressed cancer driver genes in regions with somatic copy number gains with high accuracy. We could even determine the expressed isoform of genes with several TSSs as confirmed by RNA-Seq of the matching primary tumor. Our analyses provide functional information about the cells releasing their DNA into the circulation.

---

Cell-free DNA (cfDNA) from plasma is an intensively investigated biomarker. In the field of oncology, numerous publications have demonstrated that analyses of cancer cell-derived DNA in the circulation, referred to as circulating tumor DNA (ctDNA), can be used to track tumor dynamics in real time ^1^^-^^5^. cfDNA fragments have been associated with the release of DNA from apoptotic cells after enzymatic processing since the distribution of their lengths has a mode near 166 bp in most analyses, a size which corresponds approximately to the DNA wrapped around a nucleosome (~147 bp) plus a linker fragment (~20 bp) ^6^^-^^8^. Indeed, evidence that cfDNA reflects nucleosome footprints was recently reported ^9^.

Importantly, micrococcal nuclease (MNase) assays, in which MNase digestion is used to produce mononucleosome-bound DNA fragments to define nucleosome positions in genomes, have revealed specific nucleosome patterns at promoters, which profoundly influence gene regulation ^10^^-^^13^. In actively transcribed genes, the promoter region, i.e. the region of about 150 bp upstream of the transcriptional start site (TSS), is a nucleosome-depleted region (NDR) that facilitates access to the bulky transcriptional machinery, which is flanked by arrays of well-positioned nucleosomes ^11^^-^^13^. Furthermore, a reduction in nucleosome occupancy was found which extended up to 1 kb into the gene body, resulting in reduced frequencies of mapping reads ^11^,^13^. Inactive promoters, by contrast, exhibited neither a pronounced depletion nor strong positioning and phasing of nucleosomes ^13^.

Our investigation was hence threefold: to determine whether plasma DNA is able to reflect such expression-specific nucleosome occupancy at promoters, to assess if plasma DNA possesses the sensitivity and accuracy to predict whether genes are expressed or not, and furthermore, to determine if blood samples from patients with cancer are informative for expressed cancer driver genes. To this end, we conducted WGS of plasma DNA from 50 male and 54 female donors, 179 paired-end sequenced plasma samples and 2 patients with metastasized breast cancer and we generated altogether over ~414 Gbp of raw sequencing data and ~2.6 billion mapped plasma sequence reads. We then analyzed 426 additional plasma samples from cancer patients for their suitability for nucleosome promoter analysis.

Analyses of 179 paired-end sequenced plasma DNA samples confirmed the expected unimodal size distribution of plasma nuclear DNA fragments with a narrow range and mode at 166 bp, which differed from plasma mitochondrial DNA, in which higher-order nucleosome packaging is absent, thus leaving it more exposed to enzymatic cleavage ^14^ (Fig. 1a).

**Figure 1.**
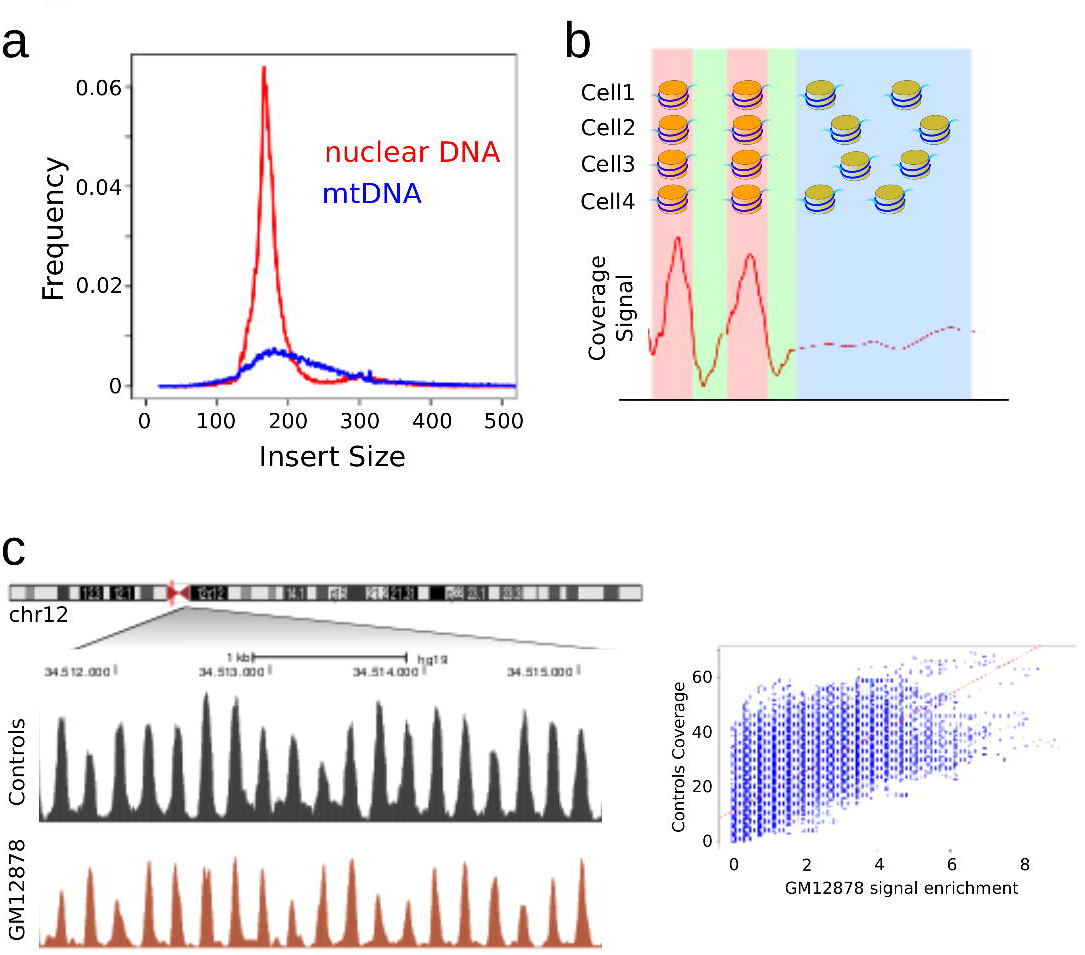
Plasma DNA fragment size and patterns of nucleosome positioning. (a) Size distribution of nuclear (red, based on ~110,000 reads) and mitochondrial (blue; based on ~53,000 reads) plasma DNA fragments. (b) Nuclear chromatin digested by micrococcal nuclease (MNase) is enriched of DNA protected by nucleosomes (dark blue) and depleted of linker regions in between (light blue). In MNase assays, regions with “perfectly positioned” nucleosomes result in strong peaks of reads reflecting the phasing of nucleosomes (left side), which differs from regions with less preferentially positioned nucleosomes (right side) [adapted from ^13^]. (c) Ideogram of chromosome 12 with enlargement of 12p11.1, which contains an extreme example of sequence-directed nucleosome positioning ^10^. The read depth analyses of plasma DNA fragments from female and male donors are shown in black and the MNase midpoint density maps from cell line GM12878 in red. To the right is the comparison between the plasma DNA read depth and MNase midpoint density maps demonstrating a strong correlation (Pearson: 0.709; Spearman: 0.708).

Sequencing of DNA fragments after MNase digestion has generated nucleosome maps where dyads (regions occupied by the center of a nucleosome) of “perfectly positioned” nucleosomes, i.e. sites with high nucleosome preferences, resulted in a strong peak of reads, reflecting the phasing of nucleosomes, whereas dyads of less preferentially positioned nucleosomes showed reduced peaks or none at all ^13^ (Fig. 1b). Near the centromere of chromosome 12 (12p11.1), there is a ~76 kb spanning region which contains over 400 consistently positioned nucleosomes independent of tissue type ^10^ and in this region, we compared the plasma DNA read counts from female and male donors with those derived from high-throughput single-end sequencing of MNase-digested chromatin of the cell line GM12878 (lymphoblastoid cell line from a female donor) taken from the ENCODE project (Fig. 1c). The plasma DNA read depth maps demonstrated wave-like patterns with peaks whose position showed a high correlation to those found in the MNase-maps (Fig. 1c). This suggested that our plasma DNA WGS data could be used to infer nucleosome positions in the human genome.

Next, we compared the read depth patterns at TSSs of 3,804 housekeeping genes ^15^ with 670 genes unexpressed in all tissues (fantom.gsc.riken.jp/5/) from the 104 controls samples. The patterns corresponded to those established by MNase assays ^11^^-^^13^ with depleted coverage at the TSS and oscillating periodicity upstream and downstream of the TSS and reduced frequencies of mapping reads (Fig. 2a). At promoters of inactive genes, by contrast, the coverage increased, reflecting the denser nucleosome packaging of repressed genes ^13^ (Fig. 2a). We then wanted to test genes expressed in blood and as spacing of nucleosomes differs between cell types ^13^, we used the cell line GM12878 for comparison with plasma read depth coverage from samples of healthy donors, from which the vast majority (>90%) of DNA fragments are derived from white blood cells ^16^,^17^. For the 1,000 (representing 1,334 TSS) most highly and the 1,000 (1,109 TSS) least expressed genes in blood ^18^, we observed similar coverage patterns with both publicly available GM12878 MNase data sets ^19^ and our own plasma DNA fragments (Fig. 2b-c). These plasma TSS sequence read depth maps differed depending on the expression level of genes (Fig. 2d).

**Figure 2.**
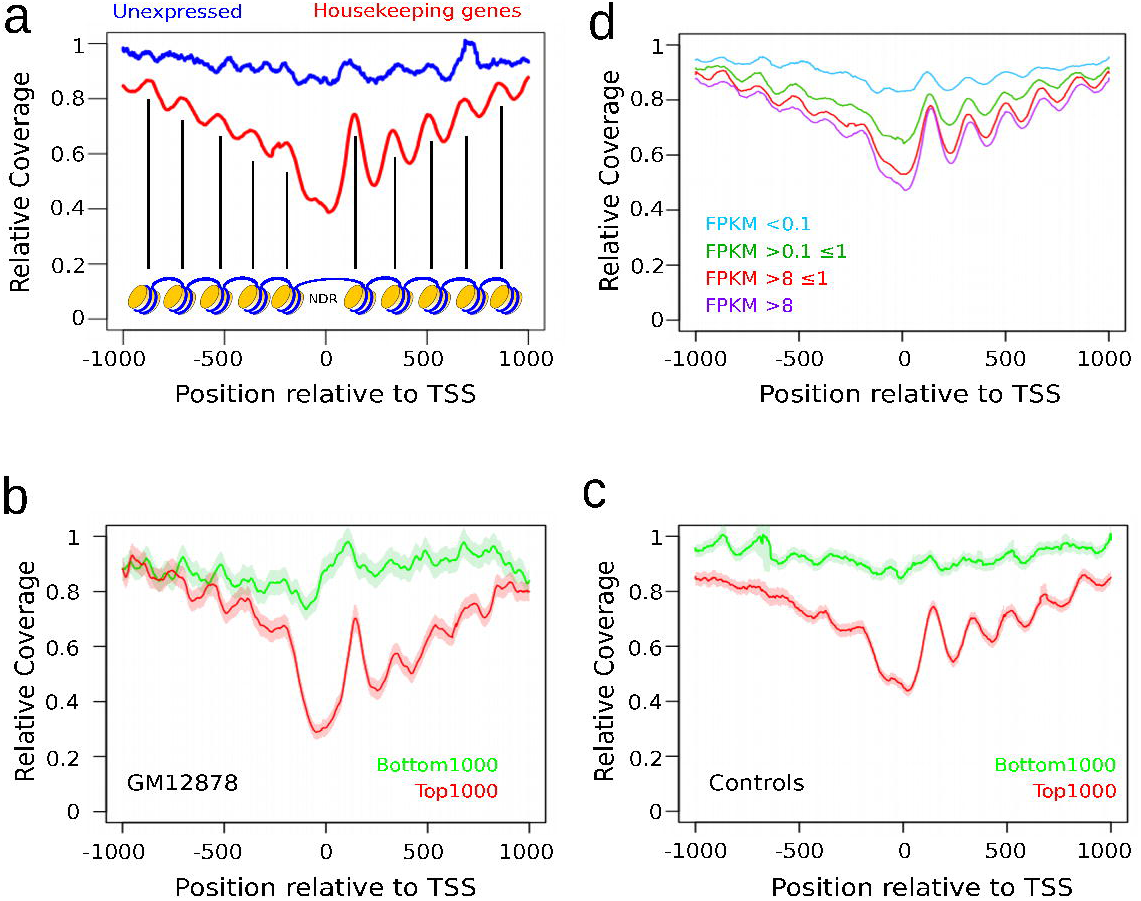
Nucleosome positioning at transcription start sites. (a) Sequencing coverage at promoter sites in housekeeping (red) and unexpressed (blue) genes generated with plasma samples from 104 donors. The coverage pattern reflects the nucleosome organization: At transcription start, nucleosomes are removed to create a nucleosome-depleted region (NDR) over the promoter, allowing transcription factors to bind ^11^^-^^13^. The reduction in nucleosome occupancy of the expressed housekeeping genes also resulted in a decreased coverage (x-axis: distance from the TSS; y-axis: relative coverage reflecting nucleosome dyads). (b) MNase midpoint density maps of GM12878 for the 1,000 (representing 1,334 TSS) most highly (red) and 1,000 least (representing 1,109 TSS) (green) expressed genes in blood based on published plasma RNA-Seq data ^18^. (c) Plasma DNA read depth maps of promoter regions of the same genes as in Fig. 2b (red: 1,000 most highly expressed genes; green: 1,000 least expressed genes). (d) Plasma DNA read depth patterns at promoters of differently expressed genes (i.e. FPKM [fragments per kilobase of mature transcript per million mapped reads] >8 (purple); FPKM >1 and ≤8 (red); FPKM >0.1 and ≤1 (green); FPKM ≤0.1 (blue)).

To distinguish between expressed and silent genes based on plasma coverage characteristics, we conducted multiple tests which resulted in the identification of two discrete regions. The first region is based on the aforementioned nucleosome occupancy reduction ±1,000 bp around the TSSs ^13^ [“2K-TSS coverage”], which we had confirmed (Fig. 2). The second region is the most frequent position of the NDR, which we mapped to −150 bp to +50 bp with respect to the TSS [“NDR coverage”] (Supplementary Fig. 1). The read depth coverage of both regions was normalized by the relative copy number so that copy number alterations, which are frequently observed in plasma samples of patients with cancer ^20^, did not affect the evaluation (Material & Methods).

Kernel density estimation of the read-depth coverage of these two regions for the 1,000 highest and least expressed genes resulted in two separate clusters (Fig. 3a). To test whether these two clusters correspond to differently expressed genes, we classified them by employing support vector machines (SVM) (Material & Methods) which allowed us to predict the expression status of the 100 most highly and least expressed genes ^18^ with a sensitivity and an accuracy of 0.91 each (Fig. 3b). Even for the 1,000 and 5,000 most highly and least expressed genes, the sensitivity was still 0.83 and 0.80, respectively, and the accuracy was 0.84 and 0.77, respectively (Fig. 3b; Supplementary Fig. 2). Accordingly, the genes in the two clusters were statistically significantly differently expressed (Mann-Whitney U Test, *p*<2.2e-16) (Fig. 3c). As examples, we illustrate the different promoter coverage patterns of the genes *NCL* and *GABRR3* (Fig. 3d). We conclude that the nucleosome protection pattern of plasma DNA allows for the distinction of expressed and silent genes with high sensitivity and accuracy and, furthermore, that the plasma of healthy donors reflects the expression signature of hematopoietic cells.

**Figure 3.**
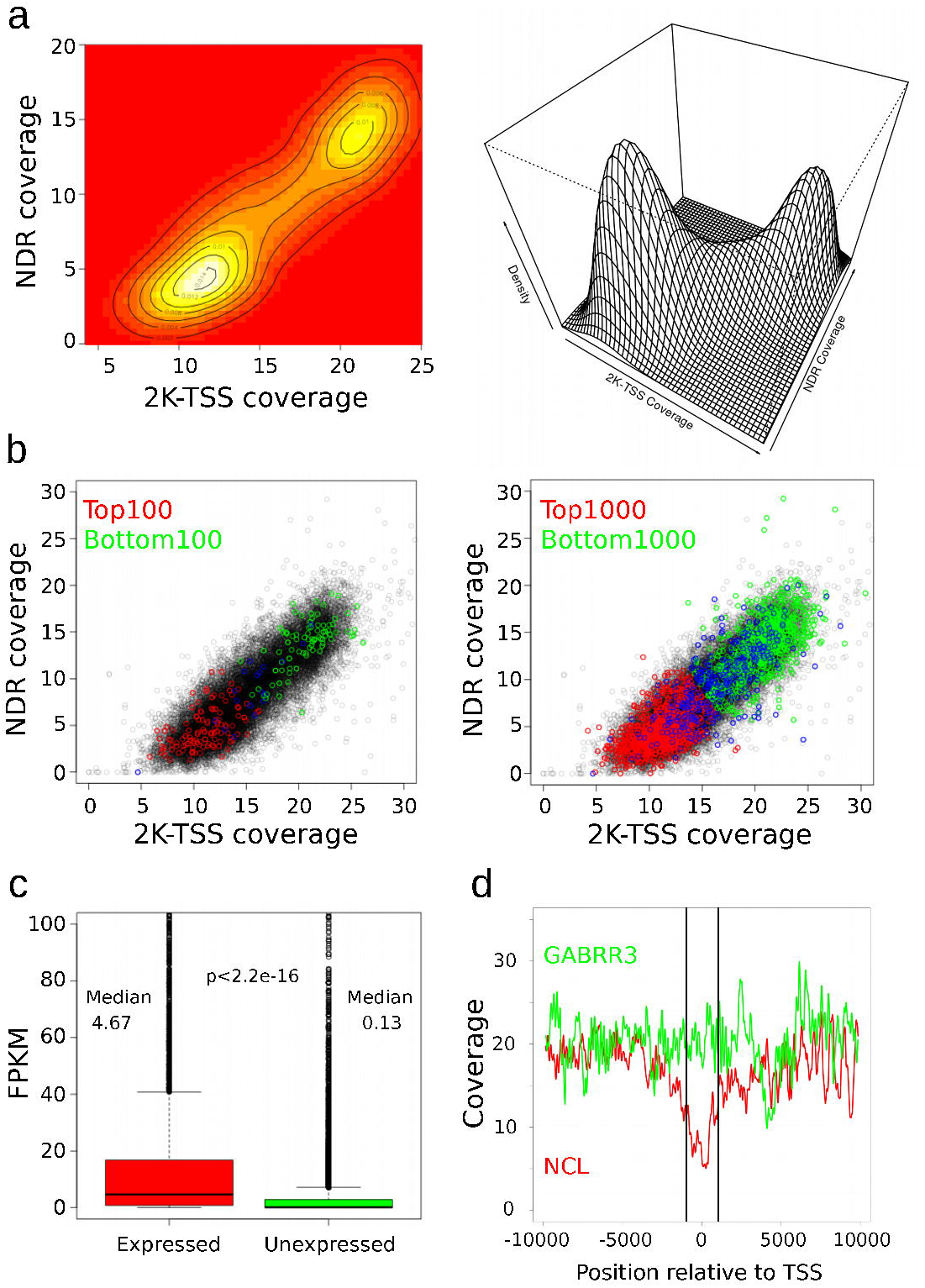
Classification of expressed and silent genes by plasma DNA read depth analyses. (a) Kernel density estimation revealed two separate gene clusters based on the normalized coverage patterns at the 2K-TSS and NDR regions. (b) Support vector machines (SVM) classification based on normalized 2K-TSS and NDR coverages for the 100 (left) and 1,000 (right) most highly and least expressed genes. Red and green circles indicate genes correctly predicted to be expressed or unexpressed, respectively, whereas blue circles represent incorrectly predicted genes. (c) The difference in FPKMs (fragments per kilobase of mature transcript per million mapped reads) for genes predicted to be expressed (n=11,345) or unexpressed (n=9,156) is statistically highly significant (Mann-Whitney U test; two-sided; median_expressed_: 4.67, sd_expressed_: 675.3, median_unexpressed_: 0.13, sd_unexpressed_: 97.0, *p*<2.2e^−16^). (d) Illustration of exemplary promoter coverage patterns of genes *NCL* (red) and *GABRR3* (green), which are expressed with mean FPKMs of 2,000 and <0.5, respectively.

We then investigated whether plasma DNA from patients with cancer would allow us to draw conclusions regarding the expression of genes in their primary tumor. Due to the inevitable heterogeneity of these plasma samples (i.e. mixture of DNA released from tumor and hematopoietic cells in various proportions), we conducted *in silico* dilution simulations to establish the resolution limits, which revealed that for this application, ≥75% of all DNA fragments of a given TSS need to be released by tumor cells in order to infer the expression status (Fig. 4a). For our proof-of-concept studies, we analyzed matching and synchronously obtained primary tumors from two metastasized breast cancer cases (B7, B13) in addition to the plasma DNA by whole-genome sequencing (Supplementary Fig. 3) and RNA-Seq (Fig. 4b). We sequenced the plasma DNA with high coverage (B7: ~411 million reads; ~8.2x; B13: ~455 million reads; ~9.1x) and calculated copy number alterations ^21^^-^^23^ (Fig. 4c). As expected, the chromosome 12p11.1 nucleosome array (Supplementary Fig. 4) and promoter read-depth differences between unexpressed and housekeeping genes (Supplementary Fig. 5) could again be established with these samples. We estimated tumor purity directly from observed relative copy profiles ^24^ and found overall ctDNA allele frequencies (AFs) of ~45% and ~72% in B7 and in B13, respectively (Fig. 4d), which are ctDNA AFs common in metastatic disease ^25^. However, the actual ctDNA AF of a distinct region additionally depends on its copy number, as amplified regions are relatively enriched for ctDNA. Therefore, we calculated regional ctDNA AFs based on the overall ctDNA AF and the log2-ratio of the respective region (Fig. 4e), which suggested that accurate gene expression predictions should at least be possible for chromosome 1q and the amplified regions on chromosomes 11q, 16p, and 19p in B7, whereas all over-represented regions should be suitable in B13.

**Figure 4.**
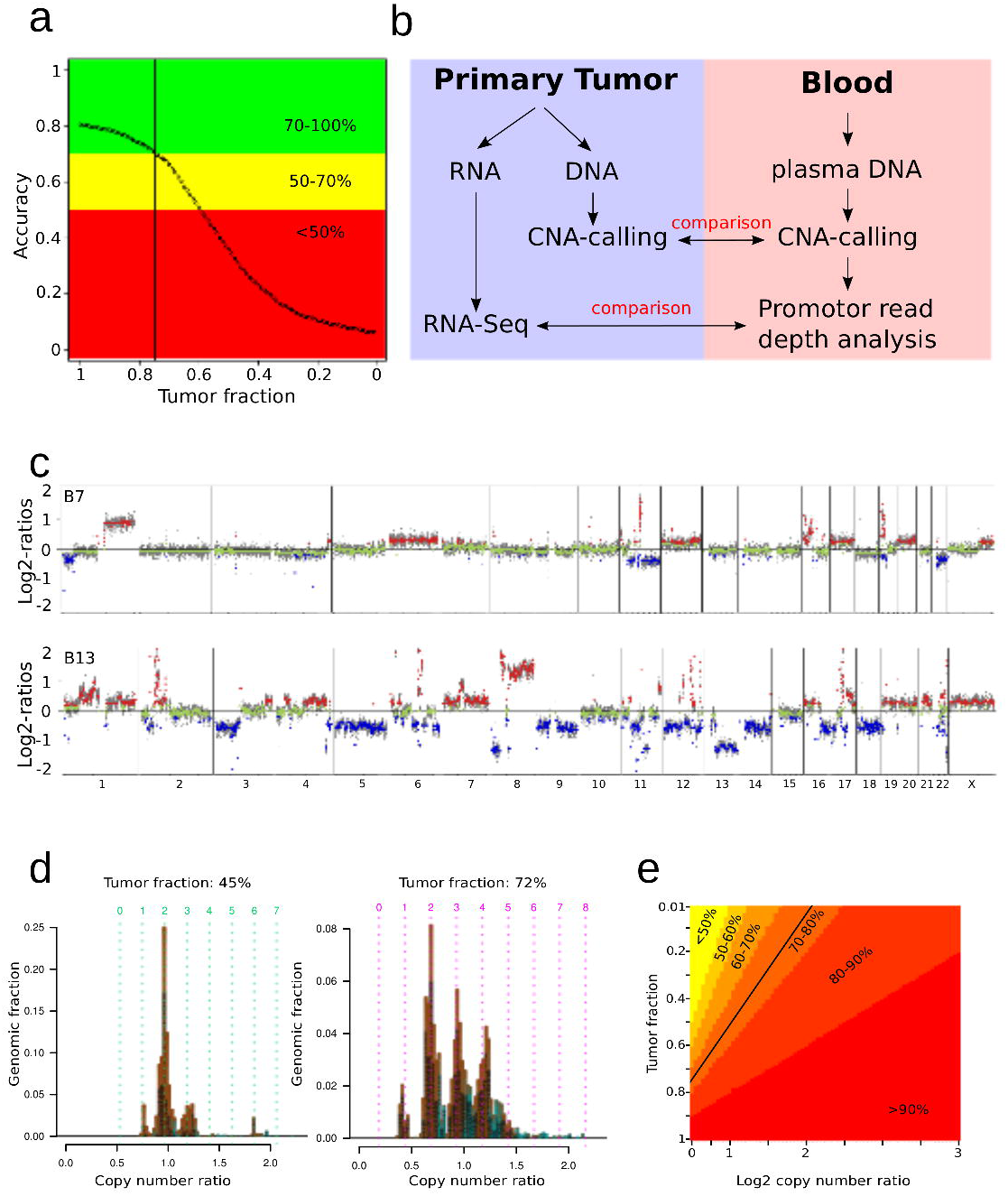
Procedure for predicting expressed driver cancer genes in blood. (a) Simulation of resolution limits with *in silico* dilution employing the 2K-TSS and the NDR coverage mixing the 1,000 most highly expressed genes with random parameters from the distribution of the 1,000 least expressed genes in plasma (green: accuracy of >70%; yellow: accuracy of 50-70%; red: accuracy below 50%). (b) Identification of expressed driver cancer genes in the peripheral blood: Matching primary tumor tissue was synchronously obtained with the blood samples. Copy number alterations (CNA) from both the primary tumor and plasma DNA were established for comparison. Expression patterns in the primary tumor were analyzed by RNA-Seq and correlated with the plasma DNA promoter coverage in relation to the respective copy number status. (c) Copy number profiles of two patients with breast cancer (B7 and B13) from plasma. The X-axis shows the chromosomes, the Y-axis indicates log2 copy number ratios. (d) Estimation of tumor purity and ploidy by the quantitative ABSOLUTE method, which estimates tumor purity and ploidy directly from observed relative copy profiles ^24^, for B7 (left) and B13 (right). (e) A heatmap illustrating how the regional ctDNA AFs are established in relation to the overall ctDNA AF (y-axis) and copy number (log2-ratio) (x-axis).

We identified focal amplifications as defined previously ^26^ which are frequent in breast cancer, such as amplifications of 11q13.3 (15 genes including *CCND1*) in B7 or of 8p11 (31 genes including *FGFR1*) and 17q12 (46 genes including *ERBB2*) in B13 (Fig. 4c). We compared the FPKMs of each gene predicted to be expressed with those predicted to be not expressed in these amplicons for B7 and B13 and observed statistically highly significant differences (Fig. 5a). We then analyzed the 100 most highly expressed genes as determined by RNA-Seq from the primary tumor from chromosome 1q in B7 and from chromosome 8p11-qter in B13 and found that 86.1% and 88.1%, respectively, were correctly classified in the expressed cluster (Fig. 5b). When we extended these analyses to the 100 most highly expressed genes in all gained regions of B13 (i.e. regions with log2-ratio>0.2; corresponding to ~1 Gbp), 78.0% were correctly classified. To provide more detailed examples for single genes, we sought to determine which isoforms were expressed in B13 for two highly relevant cancer genes, i.e. *ERBB2*, which is an important biomarker for treatment decisions involving the monoclonal antibody trastuzumab ^27^, and *FGFR1*, a potential target for fibroblast growth factor receptor (FGFR) inhibitors, which are currently in development ^28^. *ERBB2* had the promoter coverage of an expressed gene (Fig. 5c). For its two isoforms (NM_004448 and NM_001005862), we calculated the differences in the distances between the 2K-TSS and NDR coverage in blood from cancer patients and those from healthy controls (Material & Methods). This calculation predicted NM_004448 as the highly expressed isoform in the primary tumor (Fig. 5d), which was indeed confirmed by RNA-Seq [NM_004448: 11.4 FPKM; NM_001005862: 4.4 FPKM]. In fact, of 20,816 TSSs, 98.5% had a lower Euclidean distance than the second isoform of *ERBB2*. Using the same approach, we analyzed *FGFR1*, which has 2 TSSs but 9 isoforms, and could show that TSS2 should be more highly expressed than TSS1, which correlated again with the RNA-Seq data (TSS1: 3 isoforms: chr8:38325363; Sum FPKM: 6.5; TSS2: 6 isoforms: chr8:38326352; Sum FPKM: 3.0) (Fig. 5e). This prompted us to analyze every gene in the focal amplifications with several TSSs. Of the 93 genes, 8 had more than 1 TSS which gave rise to isoforms with at least 2 FPKM difference (including *ERBB2* and *FGFR1*). We were able to verify the highest expressed isoform in 7 of these again using Euclidean distances from controls to tumor samples of the same TSS (Fig. 5f). As our evaluations depend on the ctDNA AF and the log2-ratios of respective regions, we wanted to test whether this approach is broadly applicable. To this end, we analyzed 426 plasma samples from patients with metastasized cancer (colon=128; prostate=139; breast=125; lung=31; other tumor entities=3) and calculated the overall ^24^ and regional ctDNA AFs, which revealed that 220 (51.6%) of these samples had at least 100 bins (>5.6 Mbp) suitable for promoter read depth analysis. Certain regions such as high-level amplifications will almost always be amenable to our analyses, which is important as they frequently contain important cancer driver genes ^26^,^29^,^30^. Hence, in more than half of all cancer patients with common AFs of ctDNA in metastatic disease ^25^, we should be able to predict the expression of cancer genes in regions relatively enriched with ctDNA. However, our approach will likely not be applicable in minimal residual disease scenarios.

**Figure 5.**
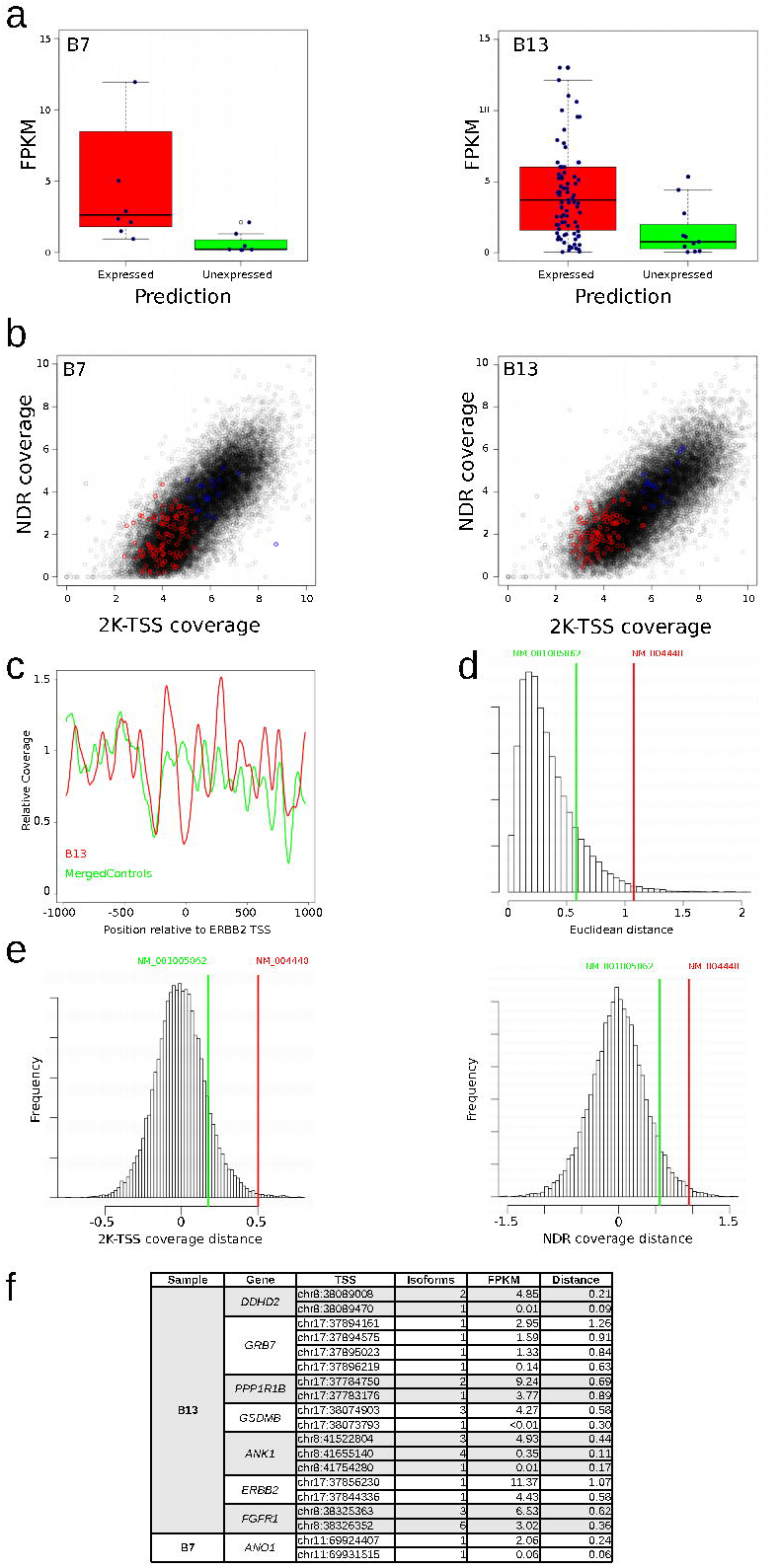
Identification of expressed driver cancer genes in the peripheral blood. (a) FPKMs for genes predicted to be expressed or not expressed in focal amplifications of 11q13.3 (15 TSS in 15 genes including *CCND1*; n_expressed_ = 8; n_unexpressed_=7) in B7 (left) and in both 8p11 (39 TSSs in 31 genes including *FGFR1*) and 17q12 (59 TSSs in 46 genes including *ERBB2*) (right) in B13 (n_expressed_ = 87; n_unexpressed_=11). Blue dots represent genes located in the amplicons. Outliers including *CCND1* (FPKM of 50 in B7) and *ERBB2* (FPKM of 15 in B13) are not shown due to scaling. The differences were statistically highly significant (One-sided Mann Whitney U tests; B7: mean_expressed_: 9.7 FPKM, sd_expressed_: 17.0, mean_unexpressed_: 0.7 FPKM, sd_unexpressed_: 0.8, *p*=0.003; B13: mean_expressed_: 5.7 FPKM, sd_expressed_: 9.7, mean_unexpressed_: 1.5 FPKM; sd_unexpressed_: 1.8, *p*=0.001) (b) Classification accuracy for the 100 most highly expressed genes in chromosomes 1q in B7 and 8p11-8qter in B13. (c) The different promoter coverage of *ERBB2* (mean of two isoforms) in B13 and in control samples. (d) *ERBB2* has two isoforms (NM_001005862 and NM_004448) and calculation of the differences in the Euclidean distances of the 2K-TSS and NDR coverage in blood from patients with cancer to those from healthy controls established that isoform NM_004448 was highly expressed in patient’s B13’s tumor. (e) Separate distances of the 2K-TSS (left panel) and NDR coverage (right panel) confirms that isoform NM_004448 was highly expressed in patient B13’s tumor. (f) The TSS giving rise to higher expressed isoforms of genes with several TSSs was identified in 7 of 8 genes within focal amplifications in B13 and B7.

Our study suggests that read depth analyses of plasma DNA can reveal functional data such as the expression status of genes due to the nucleosome occupancy pattern at promoter regions and, furthermore, that even the expression status of cancer-related genes can be deduced from the blood of patients with cancer. However, despite the high accuracy which was achieved for the prediction of expressed genes, there are factors hampering these analyses, such as the inevitable heterogeneity of plasma DNA. Furthermore, a nucleosome-deprived, regulatory factor-accessible state occurs not only in expressed, but also in paused genes, as genes with elongating Pol II or with poised Pol II exhibited a similar pattern of nucleosome phasing as expressed genes ^11^,^31^. Vice versa, transcription is not always associated with chromatin reorganization ^12^. Recently, nucleosome spacing was used to determine cfDNA tissues-of-origin ^9^. Using plasma samples from 5 individuals with cancer, these authors found correlations to the correct non-hematopoietic cell sources in three of the five tested cases based on the nucleosome footprints ^9^. This added to a previous publication, in which genome-wide bisulfite sequencing of plasma DNA with reference to methylation profiles of different tissues was used to identify the tissue contributors of the circulating DNA pool ^17^. In contrast, to the best of our knowledge, our study is the first using plasma DNA nucleosome patterns at promoter regions for the establishment of the expression status of genes. Despite the aforementioned caveats, our strategy and approach will pave the way for novel biological applications. For example, it would be interesting to test our approach in other conditions associated with increased cfDNA levels due to tissue damage, such as myocardial infarction, stroke, or autoimmune disorders. In cancer, we demonstrate that the majority of metastasized cases are amenable for non-invasive gene expression promoter read depth analysis. Due to the increasing application of molecularly driven therapeutics, which rely on accurate and timely measurements of critical biomarkers, our results are of utmost significance, as they provide a new view on the genomes of the cells which release their DNA into the circulation. Furthermore, our approach is applicable during a disease stage at which most clinical studies in oncology are conducted. This significantly expands upon the currently existing options for cfDNA analysis.

## Acknowledgements

This work was supported by CANCER-ID, a project funded by the Innovative Medicines Joint Undertaking (IMI JU). We are grateful to Samantha Perakis for the critical reading and editing of this manuscript.

## Author contributions

P.U. and M.R.S. designed the study. M.A. and R.G. performed the experiments. P.U., G.G.T., J.B.G., E.H., and M.R.S. analyzed data. E.P. provided clinical samples and clinical information. S.W.J. and L.A. performed pathologic analyses. K.K. conducted RNA-seq. P.U., E.H., and M.R.S. supervised the study. P.U., J.B.G., E.H., and M.R.S. wrote the manuscript. All authors revised the manuscript.

## Material & Methods

### Patients

The study was approved by the Ethics Committee of the Medical University of Graz (approval numbers 21-227 ex 09/10 and 21-228 ex 09/10), conducted according to the Declaration of Helsinki and written informed consent was obtained from all patients.

### Plasma DNA preparation

Plasma DNA was prepared using the QIAamp DNA Blood Mini Kit (Qiagen, Hilden, Germany) as previously described. Samples selected for sequencing library construction were analyzed on the Bioanalyzer instrument (Agilent Technologies, Santa Clara, CA, USA) to observe the plasma DNA size distribution.

### Sequencing

Shotgun libraries of plasma DNA and tumor DNA were prepared using the TruSeq DNA Nano library preparation kit by Illumina (Illumina, San Diego, CA, USA) with a starting amount of 5-10ng according to the protocol. However, due to the low DNA input, we increased the amount of PCR cycles to 25. Furthermore, the fragmentation step was omitted due to the degradation of plasma DNA. Libraries were sequenced on the Illumina MiSeq and NextSeq sequencers. All sequencing raw data were deposited at the European Genome-phenome Archive (EGA, http://www.ebi.ac.uk/ega/), which is hosted by the EBI, under the accession number EGAS00001001754.

### Paired-end sequencing data preparation

Paired-end reads of 179 plasma DNA samples were aligned with bwa ^32^ backtrack to the human hg19 genome. Resulting BAM files were merged using samtools ^33^ and alignments to the mitochondrial genome were extracted. Nuclear alignments were downsampled and insert sizes of both BAM files were analyzed using Picard’s InsertSizeMetrics function.

### CNA analysis

Raw reads of the two breast cancer samples and the merged controls were aligned to the human hg19 genome using bwa ^32^ where the pseudo-autosomal region of the Y-chromosome was masked. PCR duplicates were removed and reads were counted in 50,000 genome bins, each containing the same amount of mappable positions (approximately 56kbp). Raw read counts were normalized by the median bin count and GC correction was done using LOWESS smoothing. Furthermore, corrected read counts were normalized by mean bin counts of non-cancer controls and segmented using both CBS and GLAD provided by the CGHweb framework^34^.

### Tumor fraction estimation

The tumor fraction of the two breast cancer samples was estimated by applying ABSOLUTE ^24^ to the segmented log2-ratios obtained by the CNA analysis. We used the most plausible karyotype model and extracted purity values.

### Relative tumor fraction estimation

In order to estimate the relative tumor fraction of a region depending on the copy number state,
we derived the following:

The undiluted copy number (cp_i_) of a certain region i depends on the log2-ratio (lr_i_) as measured in the CNA analysis step as well as the tumor fraction (tf)

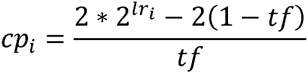

The relative tumor fraction (rtf,_i_) for this region can then be computed using the (pure) copy number and the tumor fraction again.

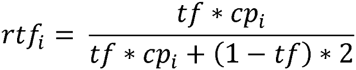

### Single-end sequencing data preparation

Raw reads (150bp) of the 104 control samples and the two breast cancer samples were trimmed from both ends to contain bases 53 to 113. These 60bp should constitute the central 60bp of a typical 166bp cfDNA fragment and should thus be exclusively associated with a nucleosome. Reads were then aligned to the human hg19 genome using bwa-mem (version 0.7.4) ^32^ and PCR duplicates were removed using the samtools rmdup ^33^ function (version 0.1.18). Aligned BAM files of controls were merged using the samtools merge function.

### Plasma RNA analysis

Gene expression values of plasma RNA analyses from microarrays were provided by Koh et al. RMA values of four healthy (non-pregnant) subjects were averaged and the 1,000 most expressed genes (Top1000) and 1,000 least expressed genes (Bottom1000) were extracted. Moreover, raw fastq files from the RNA-Seq step were downloaded from the four non-pregnant samples (SRA accessions: SRR1296080, SRR1296081, SRR1296082, SRR1296083) and analyzed as detailed below.

### RNA-Seq

RNA-Seq expression values were computed from raw data provided from ^18^ and from tumor
samples of B7 and B13. Briefly, we aligned RNA-Seq reads to the human hg19 genome using TopHat2 (v2.0.7) and calculated the gene-wide FPKM for each of the four samples using Cufflinks2 ^35^. Subsequently, FPKMs of the four samples were averaged for each gene.

### TSS profile

Coverage values around transcription start sites were extracted from aligned BAM files using the samtools depth ^33^ function and every value was normalized by the mean value of the regions: TSS-3000 to TSS-1000 and TSS+1000 and TSS+3000, respectively.

### Copy number normalized parameter extraction

Two parameters were used for the identification/prediction of genes into an expressed and unexpressed subset.

1. The coverage between TSS-1000bp and TSS+1000bp (2K-TSS coverage)
2. The coverage between TSS-150bp and TSS+50bp (NDR coverage)

For every TSS in RefSeq, parameters were extracted and divided by the relative copy number of that region identified in the CNA analysis step.

### Prediction by Support Vector Machines

In order to predict the expression status of individual genes, we used Support Vector Machines. As a training set for expressed genes, we used a random subset of 300 housekeeping genes out of 3,804 housekeeping genes which are expressed uniformly in multiple tissues ^15^ and for unexpressed genes a random subset of 300 genes out of 670 reported to be unexpressed in most tissues by the FANTOM5 project (fantom.gsc.riken.jp/5/). The remaining genes in both classes were used as the test set and their expression status predicted. Random subset selection and prediction was repeated a 1,000 times and prediction status for each TSS was recorded. We considered a gene to be expressed when the prediction consent of all the iterations was higher than 75%.

### In-silico dilution

We performed dilution simulations to test the reliability of the prediction at varying tumor fractions. To this end, we modeled the distribution of the 2K-TSS and the NDR coverage parameters of the 1000 least expressed genes in Plasma and added random numbers from these distributions to the parameters of the Top 1000 expressed genes at varying proportions.

### Isoform discrimination

Expressed isoforms were determined by calculating the distance of the two parameters between the TSS in the merged control data and the tumor patient after normalizing both parameters in both data sets. TSSs which lead to higher expression in the tumor should decrease in both the 2K-TSS and the NDR coverage when compared to the same TSS in control data.

### Code availability

Relevant code is available at https://github.com/PeterUlz/Nucleosome_ctDNA.

## References

1. Schwarzenbach, H., Hoon, D.S. & Pantel, K. Cell-free nucleic acids as biomarkers in cancer patients. Nat Rev Cancer 11, 426–37 (2011).

2. Heitzer, E., Auer, M., Ulz, P., Geigl, J.B. & Speicher, M.R. Circulating tumor cells and DNA as liquid biopsies. Genome Med 5, 73 (2013).

3. Crowley, E., Di Nicolantonio, F., Loupakis, F. & Bardelli, A. Liquid biopsy: monitoring cancer-genetics in the blood. Nat Rev Clin Oncol 10, 472–84 (2013).

4. Diaz, L.A., Jr. & Bardelli, A. Liquid biopsies: genotyping circulating tumor DNA. J Clin Oncol 32, 579–86 (2014).

5. Heitzer, E., Ulz, P. & Geigl, J.B. Circulating tumor DNA as a liquid biopsy for cancer. Clin Chem 61, 112–23 (2015).

6. Diehl, F. et al. Detection and quantification of mutations in the plasma of patients with colorectal tumors. Proc Natl Acad Sci U S A 102, 16368–73 (2005).

7. Lo, Y.M. et al. Maternal plasma DNA sequencing reveals the genome-wide genetic and mutational profile of the fetus. Sci Transl Med 2, 61ra91 (2010).

8. Ramachandran, S. & Henikoff, S. Replicating Nucleosomes. Sci Adv 1, pii: e1500587 (2015).

9. Snyder, M.W., Kircher, M., Hill, A.J., Daza, R.M. & Shendure, J. Cell-free DNA Comprises an In Vivo Nucleosome Footprint that Informs Its Tissues-Of-Origin. Cell 164, 57–68 (2016).

10. Gaffney, D.J. et al. Controls of nucleosome positioning in the human genome. PLoS Genet 8, e1003036 (2012).

11. Schones, D.E. et al. Dynamic regulation of nucleosome positioning in the human genome. Cell 132, 887–98 (2008).

12. Venkatesh, S. & Workman, J.L. Histone exchange, chromatin structure and the regulation of transcription. Nat Rev Mol Cell Biol 16, 178–89 (2015).

13. Valouev, A. et al. Determinants of nucleosome organization in primary human cells. Nature 474, 516–20 (2011).

14. Chandrananda, D., Thorne, N.P. & Bahlo, M. High-resolution characterization of sequence signatures due to non-random cleavage of cell-free DNA. BMC Med Genomics 8, 29 (2015).

15. Eisenberg, E. & Levanon, E.Y. Human housekeeping genes, revisited. Trends Genet 29, 569–74 (2013).

16. Lui, Y.Y. et al. Predominant hematopoietic origin of cell-free DNA in plasma and serum after sex-mismatched bone marrow transplantation. Clin Chem 48, 421–7 (2002).

17. Sun, K. et al. Plasma DNA tissue mapping by genome-wide methylation sequencing for noninvasive prenatal, cancer, and transplantation assessments. Proc Natl Acad Sci U S A 112, E5503–12 (2015).

18. Koh, W. et al. Noninvasive in vivo monitoring of tissue-specific global gene expression in humans. Proc Natl Acad Sci U S A 111, 7361–6 (2014).

19. Encode Project Consortium. An integrated encyclopedia of DNA elements in the human genome. Nature 489, 57–74 (2012).

20. Heitzer, E., Ulz, P., Geigl, J.B. & Speicher, M.R. Non-invasive detection of genome-wide somatic copy number alterations by liquid biopsies. Mol Oncol 10, 494–502 (2016).

21. Heidary, M. et al. The dynamic range of circulating tumor DNA in metastatic breast cancer. Breast Cancer Res 16, 421 (2014).

22. Heitzer, E. et al. Tumor-associated copy number changes in the circulation of patients with prostate cancer identified through whole-genome sequencing. Genome Med 5, 30 (2013).

23. Mohan, S. et al. Changes in Colorectal Carcinoma Genomes under Anti-EGFR Therapy Identified by Whole-Genome Plasma DNA Sequencing. PLoS Genet 10, e1004271 (2014).

24. Carter, S.L. et al. Absolute quantification of somatic DNA alterations in human cancer. Nat Biotechnol 30, 413–21 (2012).

25. Bettegowda, C. et al. Detection of circulating tumor DNA in early-and late-stage human malignancies. Sci Transl Med 6, 224ra24 (2014).

26. Ulz, P., Heitzer, E. & Speicher, M.R. Co-occurrence of MYC amplification and TP53 mutations in human cancer. Nat Genet 48, 104–6 (2016).

27. Giordano, S.H. et al. Systemic therapy for patients with advanced human epidermal growth factor receptor 2-positive breast cancer: American Society of Clinical Oncology clinical practice guideline. J Clin Oncol 32, 2078–99 (2014).

28. Helsten, T. et al. The FGFR Landscape in Cancer: Analysis of 4,853 Tumors by Next-Generation Sequencing. Clin Cancer Res 22, 259–67 (2016).

29. Beroukhim, R. et al. The landscape of somatic copy-number alteration across human cancers. Nature 463, 899–905 (2010).

30. Vogelstein, B. et al. Cancer genome landscapes. Science 339, 1546–58 (2013).

31. Adelman, K. & Lis, J.T. Promoter-proximal pausing of RNA polymerase II: emerging roles in metazoans. Nat Rev Genet 13, 720–31 (2012).

32. Li, H. & Durbin, R. Fast and accurate short read alignment with Burrows-Wheeler transform. Bioinformatics 25, 1754–60 (2009).

33. Li, H. et al. The Sequence Alignment/Map format and SAMtools. Bioinformatics 25, 2078–9 (2009).

34. Lai, W., Choudhary, V. & Park, P.J. CGHweb: a tool for comparing DNA copy number segmentations from multiple algorithms. Bioinformatics 24, 1014–5 (2008).

35. Trapnell, C. et al. Differential gene and transcript expression analysis of RNA-seq experiments with TopHat and Cufflinks. Nat Protoc 7, 562–78 (2012).

